# Developmental trajectories of the default mode, executive control, and salience networks from the third trimester through the newborn period

**DOI:** 10.1101/2022.09.27.509687

**Authors:** Dustin Scheinost, Joseph Chang, Emma Brennan-Wydra, Cheryl Lacadie, R. Todd Constable, Katarzyna Chawarska, Laura R. Ment

## Abstract

Social cognition is critical to early learning. Functional imaging studies in adults and older children suggest the involvement of the default mode (DMN), executive control (ECN), and salience (SAL) networks in social cognition. These networks are vulnerable to environmental insults, and abnormalities of intra- and inter-network connectivity of the three are emerging as biomarkers of neurobehavioral disorders. However, the developmental trajectories of the DMN, ECN, and SAL across the third trimester of gestation and perinatal transition remain largely unknown. Employing resting-state functional MRI studies at 30-32, 34-36, and 40-44 weeks postmenstrual age (PMA), we tested the hypothesis that both intra- and inter-network functional connectivity in the DMN, ECN, and SAL develop across the 30-46 weeks PMA time interval in a longitudinal/cross-sectional sample of 84 fetuses and neonates. A secondary analysis addressed the impact of maternal mental health assessed at 28 weeks PMA on tri-network development from 30-46 weeks PMA. The DMN, ECN, and SAL develop across the third trimester of gestation and the first postnatal month. At the intra-network level, significant increases occurred between 36 to 44 weeks PMA for all three, with network strength values significantly different from 0 beginning at 40 weeks PMA for all. Functional connectivity increased less rapidly in the DMN than in the ECN and SAL networks, suggesting slower maturation of the network subserving social interactions. In contrast, significant inter-network DMN – ECN connectivity greater than 0 was found from 36 weeks PMA through the first postnatal month, suggesting the emergence of inter-network functional connectivity in the fetal brain. Finally, higher maternal mental health symptoms measured at the beginning of the third trimester negatively affected the developmental trajectory of the SAL network across the critical time interval of 30 weeks to 44 weeks PMA. Together, these data provide a framework to compare fetuses and neonates at risk for neurobehavioral disorders and assess the impact of the environment on the developing brain.

## INTRODUCTION

Social cognition—the neural process by which one processes, stores, and applies information about other people and social situations—is critical to early learning and requires the synchronous engagement of multiple distinct information-processing networks across the developing brain.^1^ Imaging studies in adults and older children suggest the involvement of three critical networks and their interconnections in social cognition: the default mode, executive control, and salience networks.^2-4^ Although neuroimaging studies demonstrate the functional organization of the fetal connectome across the second and third trimesters of gestation,^5-9^ in typically developing fetuses and neonates, these networks and their relationships are just beginning to be explored.

The default mode, executive control, and salience networks are called the “tri-network.” The default mode network (DMN), composed of the medial prefrontal cortex, posterior cingulate cortex, and the bilateral angular gyri, subserves wakeful rest and social interactions. Whereas the executive control network (ECN), composed of the bilateral dorsolateral prefrontal cortex and intra-parietal sulci,^3,10^ maintains and manipulates information in working memory and is responsible for the decision-making required in goal-based behavior. The salience network (SAL) comprises the bilateral anterior insula and anterior cingulate cortex (ACC) and maps necessary external inputs and internal cerebral events. The SAL is proposed to initiate dynamic switching between the DMN and ECN.^2,3,11^

Although adult-like topography has been reported in primary networks, including sensorimotor, visual, and auditory cortices, at term equivalent age,^12-14^ higher-order networks, including the DMN, ECN, and SAL, are thought to be topologically incomplete and isolated from other networks at the same age,^12,13,15-18^ showing significant growth in topology and synchronization during the first three postnatal months of life. These changes are followed by less consistent growth in later months, with the components of the tri-network reaching more mature characteristics by the end of the first postnatal year.^13,18-21^ Similarly, marked changes occur in the strength of inter-network communication. Integration between the ECN and DMN increases significantly across the first postnatal year, while that between the DMN and SAL and the SAL and ECN decreases during the same time interval, suggesting progressive segregation between the two networks during postnatal development.^17^ Finally, these networks are vulnerable to environmental insults in the fetal and newborn periods (such as prenatal stress exposure), which putatively lead to an increased risk of adverse developmental outcomes. ^22-29^ However, no studies address their integration and synchronization spanning the important fetal to neonatal transition in normative development.^8^

To address this significant knowledge gap,^30-37^ we tested the hypothesis that both intra- and inter-network functional connectivity in the DMN, ECN, and SAL develop across the third trimester of gestation and the first four postnatal weeks in a longitudinal and cross-sectional sample of 84 fetuses and neonates scanned between 30 weeks–46 weeks PMA. To the best of our knowledge, these data demonstrate for the first time both the developmental trajectory of the DMN, ECN, and SAL across this critical period and prenatal inter-network connectivity among these core components of social cognition in the developing brain.

## METHODS

This work includes longitudinal and cross-sectional imaging data from two Yale School of Medicine studies and an open-source dataset obtained from the Developing Human Connectome Project (dHCP).^12^ The Yale University institutional review board approved all studies. Parent(s) provided written consent. The Developing Human Connectome (dHCP) project was reviewed and approved by the UK National Research Ethics Authority. DHCP study investigators obtained written informed parental consent. An overview of the composition of the study cohorts is presented in Table 1.

**Table 1.** Sample Characteristics

|  | Fetal-neonatal Cohort | Neonatal Cohort | dHCP | Comparison p-value |
| --- | --- | --- | --- | --- |
| Number | 29 | 31 | 24 |  |
| Number females (%) | 13 (44.8) | 19 (61.2) | 9 (37.5) | .188 |
| Birth weight (g) | 3646 +/- 640 | 3510 +/- 377 | 3438 +/- 423 | .296 |
| PMA at birth (wks) | 39.4 +/- 1.5 | 39.9 +/- 0.8 | 39.4 +/- 1.6 | .232 |
| Number SGA (%) | 0 (0.0) | 0 (0.0) | 0 (0.0) | n/a |
| Race |  |  |  | .403* |
| Asian | 1 (3.4) | 3 (9.7) | 0 (0.0) |  |
| Black – African American | 4 (13.8) | 1 (3.2) | 0 (0.0) |  |
| Native American | 0 (0.0) | 0 (0.0) | 0 (0.0) |  |
| White | 19 (65.5) | 21 (67.7) | 0 (0.0) |  |
| More than 1 race | 5 (17.2) | 6 (19.4) | 0 (0.0) |  |
| Unknown/not reported | 0 (0.0) | 0 (0.0) | 38 (100.0) |  |
| Ethnicity |  |  |  | .505* |
| Hispanic – Latinx | 5 (17.2) | 4 (12.9) | 0 (0.0) |  |
| Unknown/not reported | 1 (3.4) | 0 (0.0) | 38 (100.0) |  |
| PMA at scan (wks) |  |  |  |  |
| Scan 1 (F1) | 31.2 +/- 0.7 | N/A | N/A |  |
| Scan 2 (F2) | 35.3 +/- 0.8 | N/A | N/A |  |
| Scan 3 (NN) | 43.4 +/- 1.3 | 44.4 +/- 1.3 | 40.2 +/- 2.1 | **<.001** |
| Maternal education |  |  |  | .845* |
| < High school | 0 (0.0) | 0 (0.0) | 0 (0.0) |  |
| High school graduate | 0 (0.0) | 1 (3.2) | 0 (0.0) |  |
| Some college | 2 (6.9) | 1 (3.2) | 0 (0.0) |  |
| College graduate | 25 (86.2) | 27 (87.1) | 0 (0.0) |  |
| Unknown/not reported | 2 (6.9) | 2 (6.5) | 38 (100.0) |  |
| PMA=Postmenstrual age<br>Values are +/- SD<br>Comparison p-value represents significance level of comparison between cohorts: ANOVA for continuous variables (e.g., birth weight) and chi-squared tests for categorical variables (e.g., race). **Bold text** indicates p<.05. * p-values do not include dHCP cohort (data not available). |  |  |  |  |

### Participants

#### Yale Fetal to Neonatal Cohort

The Yale cohort consists of longitudinal data collected from the third trimester to the neonatal period as part of the Yale Autism Center of Excellence Program Project. Between April 09, 2018, and April 13, 2022, 33 pregnant women were recruited for scanning at 30–33 weeks and 34–36 weeks postmenstrual age (PMA). Three did not have usable data from either fetal scan (F1 or F2), and one fetus-to-neonate participant was small for gestational age. Table 1 shows the sample characteristics for the 29 fetuses with usable data and appropriate for gestational age status. Mothers who participated in the fetal scans were invited to have their infants participate in the neonatal functional connectivity protocol with repeat MRI within the first six weeks of life. Usable longitudinal data were available for all 29 maternal-infant dyads, with at least two out of three (two fetal and one neonatal) scans per participant. Out of 29 participants, 48.3% were scanned before the COVID-19 pandemic, with the remaining 51.7% scanned during the pandemic. Inclusion criteria for fetuses included pregnant women with no family history of autism spectrum disorder (ASD) who met the following criteria: 1) 26–28 weeks gestational age; 2) estimated fetal weight, femur length, and bi-parietal diameter appropriate for gestational age; 3) no known chromosome or structural abnormalities; 4) no brain injury on clinical ultrasound; 5) no congenital infections; 6) no known ethanol exposure since confirmation of pregnancy; 7) no known tobacco/nicotine use since confirmation of pregnancy; 8) no known illicit drug use/abuse; 9) singleton pregnancy; 10) receiving regular prenatal care; 11) maternal age 21-35 years; 12) no morbid obesity; 13) mother able to give permission. Exclusion criteria included 1) any contraindications for MRI scanning; 2) too large to fit in the MRI machine comfortably for 45 minutes; 3) a clinically ordered MRI scheduled in the future; 4) difficulty lying down and remaining still for 45–60 minutes; 5) current drug, alcohol, and tobacco/nicotine use; and 6) preterm birth (for neonates). Drug screening (Integrated E-Z Split Key Cup II, Alere San Diego, Inc, San Diego, CA 92121) and alcohol and tobacco screening tools were administered to all mothers at the time of each fetal MRI study using the Fagerstrom Test for Nicotine Dependence and the Alcohol Timeline Followback assessments ^38,39^.

#### Yale Neonatal Cohort

Thirty-five healthy term infants born between 37 and 41 weeks of PMA were scanned within the first six weeks of their participation in the Yale Autism Center of Excellence Program Project. The scans were completed between April 30, 2018 and December 13, 2021. Four infants awoke before adequate functional data could be collected; thus, usable data are available for 31 of these subjects. (Table 1) Inclusion criteria included singleton pregnancy, term birth, and appropriate for gestational age. Exclusion criteria included: 1) congenital infections; 2) non-febrile seizure disorder; 3) hearing loss; 4) visual impairment; 5) the presence of any known chromosomal abnormality; 6) prenatal exposure to illicit drugs; 7) major psychotic disorder in first degree relatives; and 8) contraindications to MRI including non-removable metal medical implants (i.e., patent ductus arteriosus clip). Like the fetal cohort, only infants with no family history of autism or other neurodevelopmental disorders are included in this analysis. Please see Table 1 for sample characteristics) *dHCP Cohort:* The data from thirty-eight term-born infants prospectively recruited as part of the dHCP, an observational, cross-sectional Open Science program approved by the UK National Research Ethics Authority, were included in these analyses. Infants were recruited from the postnatal wards and scanned at 37–43.5 weeks PMA. dHCP exclusion criteria include a history of severe compromise at birth requiring prolonged resuscitation, a diagnosed chromosomal abnormality, or any contraindication to MRI scanning. Additional exclusion criteria imposed for this analysis included the need for the infants to be appropriate for gestational age. A review of birth weight data for the 35 infants using UK-WHO growth charts for ages 0-4 years^40^ revealed that 14/38 were small for gestational age. Thus, data from the 24 dHCP neonates are included in this analysis. Please see Table 1 for sample characteristics. Notably, race, ethnicity, and maternal education data were not available for the dHCP sample.

### Maternal mental health: Impact on connectivity

Maternal mental health data were collected at the first fetal scan for subjects in the Yale longitudinal analysis. Depression symptoms were quantified using the Edinburgh Postnatal Depression Scale (EPDS; a 10-item measure of perinatal depression, yielding possible scores ranging from 0 to 30)^41^. Anxiety was measured using the State-Trait Anxiety Inventory (STAI; a 40-item instrument measuring state and trait anxiety total scores, ranging from 20 to 80)^42^. 42 Stress symptoms were indexed by the Perceived Stress Scale (PSS-14; a 14-item measure of perceived stress with scores ranging from 0 to 56).^43^ For all three measures, higher scores indicate the presence of more symptoms. The composite maternal mental health variable was obtained by scaling and centering the original four variables, conducting principal components analysis (PCA), and using the first principal component (PC1) in the subsequent analyses. PC1 explains 73.0% of the total variance in the original data using a single variable, with loadings (eigenvectors) of 0.50, 0.46, 0.51, and 0.52 for EPDS, PSS, STAI State, and STAI Trait, respectively. There is no significant difference in PC1 score between mothers who were enrolled before March 2020 (n=14) vs. after (n=15), p=.930.

### Imaging Parameters

#### Fetal-neonatal Cohort

All MRIs were performed in a natural, unmedicated state.Fetuses were studied using repeat MRI protocols completed in <60 minutes using a 3 Tesla Siemens (Erlangen, Germany) Prisma MR system and a flexible, lightweight (∼1lb) cardiac 32-channel body coil. Five functional runs were acquired (TR=1950 ms, TE=21 ms, FoV=320mm, flip angle 90°, matrix size 94×94, SAR<0.4, slice thickness 3mm, Bandwidth=2215Hz/pixel, 32 slices). Each of the five functional runs comprised 150 volumes (5.85 minutes). On average, 520 frames (range: 300 - 750) were retained for analysis or more than 17 minutes of data per participant. Follow-up neonatal MRI in this cohort occurred as part of a natural-sleep “feed and wrap” protocol ^44^. Infants were fed, bundled with multiple levels of ear protection, and immobilized in an MRI-safe vacuum swaddler. Heart rate and O_2_ saturation were continuously monitored during all scans. The same 3 Tesla Siemens (Erlangen, Germany) Prisma MR system employed for fetal imaging was also used for the neonatal data. Functional images were collected using an echo-planar image gradient echo pulse sequence (TR=2120ms, TE=22ms, FoV=260mm, matrix size=102×102, slice thickness=3mm, Flip Angle=90°, Bandwidth=2335Hz/pixel, 32 slices). Functional runs consisted of 360 volumes (6.18 minutes).

#### Neonatal Cohort

Neonatal imaging was performed on the same 3 Tesla Siemens (Erlangen, Germany) Prisma MR system using a 32-channel parallel receiver head coil and the same “feed and wrap” MRI protocol as above. We collected five functional runs, each comprised of 360 volumes. On average, 682 frames (range 285–750) were retained for analysis, and each neonate had an average of 11.5 min (SD=1.4) of usable, functional data.

dHCP cohort: Imaging was acquired at the Evelina Newborn Imaging Centre, Evelina London Children’s Hospital, using a 3 T Philips Achieva system (Philips Medical Systems). All infants were scanned without sedation in a scanner environment, including a dedicated transport system, positioning device, and a customized 32-channel receive coil with a custom-made acoustic hood. MRI-compatible ear putty and earmuffs were used to provide additional acoustic noise attenuation, and infants were fed, swaddled, and positioned in a vacuum jack prior to scanning to provide natural sleep.^12^

### Image Processing

#### Fetal connectivity preprocessing

Functional data were processed using validated fetal fMRI pipelines ^45,46^. Functional data were corrected for motion using a two-pass registration approach optimized for fetuses to correct for large and small head movements ^45^. Outlying frames were censored for data quality based on the signal-to-noise ratio within the fetal brain, the final weighted correlation value from optimization, and the frame-to-frame motion between adjacent frames. These frames were defined as frames with SNR, registration quality, or motion greater/less than one standard deviation above/below the mean values over all runs.

As previously outlined^47^, several covariates of no interest were regressed from the data, including linear and quadratic drifts, six motion parameters, the mean cerebral-spinal-fluid (CSF) signal, the mean white-matter signal, and the mean gray matter signal. The data were temporally smoothed with a zero mean unit variance Gaussian filter (approximate cutoff frequency=0.12 Hz). A gray-matter mask defined in template space was applied to the data, so only gray matter voxels were used in further calculations.

Next, to warp the network seeds from MNI space to fMRI space, a series of nonlinear registrations were calculated independently and combined into a single transform. This single transformation allows the seeds to be transformed into a single participant’s space with only one transformation, reducing interpolation error. First, the mean functional image from the motion-corrected fMRI data was registered to an age-appropriate template (i.e., 31 weeks or 34 weeks gestation) ^48^ using a low-resolution nonlinear registration. These age-appropriate fetal templates were non-linearly registered to MNI space.

#### Infant connectivity preprocessing

Functional data for infants were processed using a previously validated pipeline ^44^. Functional images were motion-corrected using SPM8. Next, images were iteratively smoothed until the smoothness of any image had a full-width half maximum of approximately 6mm using AFNI’s 3dBlurToFWHM. This iterative smoothing reduces motion-related confounds ^49^. All further analyses were performed using BioImage Suite ^50^ unless otherwise specified. Several covariates of no interest were regressed from the data, including linear and quadratic drifts, mean cerebral-spinal-fluid (CSF) signal, mean white-matter signal, and mean gray matter signal. For additional control of possible motion-related confounds, a 24-parameter motion model (including six rigid-body motion parameters, six temporal derivatives, and these terms squared) was regressed from the data. The data were temporally smoothed with a Gaussian filter (approximate cutoff frequency=0.12Hz). A canonical gray matter mask defined in common space was applied to the data, so only voxels in the gray matter were used in further calculations.

Next, to warp the network seeds from MNI space to fMRI space, a series of nonlinear registrations were calculated independently and combined into a single transformation. First, the mean functional image from the motion-corrected fMRI data was registered to a custom infant template (as in ^51^) using a previously validated algorithm ^52^. Similarly, the same algorithm registered the infant template to the MNI brain.

#### Seed intra-network connectivity

After the seeds for each of the three networks (i.e., DMN, ECN, and SAL; see Figure 1 for network seeds and Table S1 for network node coordinates; Figure 1) were warped into a single participant’s space, the time course for each seed region was then computed as the average time course across all voxels in the reference region. The time courses were correlated between each remaining seed in the network and transformed to z-values using Fisher’s transform, resulting in four functional connections per individual. Connectivity strength for each overall network was defined as the average of the individual connections within each network, resulting in three functional connections per individual

**Figure 1:**
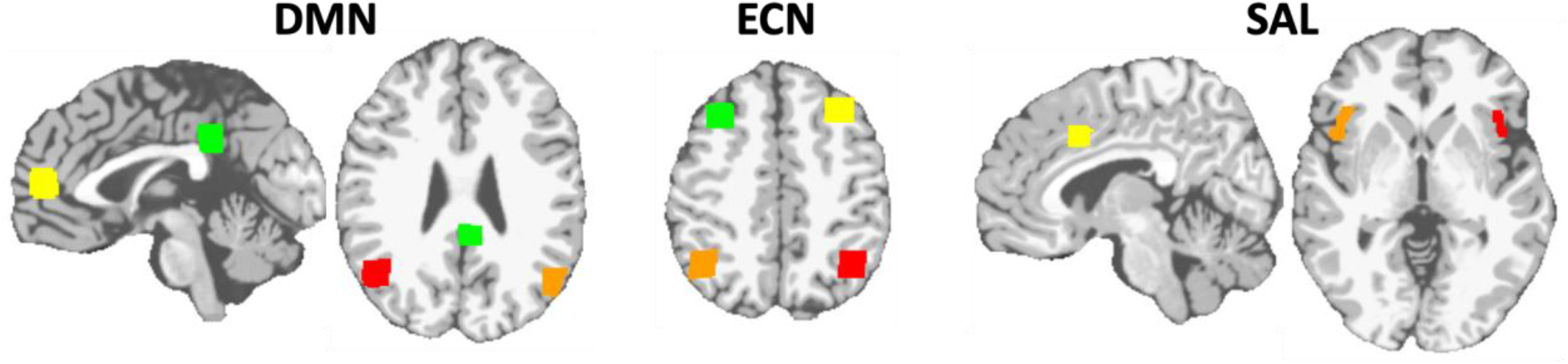
Seeds used for the DMN, ECN, and SAL networks. The DMN was defined as the PCC (green), mPFC (yellow), right angular gyrus (red), and left angular gyrus (orange). The ECN was defined as the right dlPFC (green), left dlPFC (yellow), left IPS (red), and right IPS (orange). The SAL was defined as the dACC (yellow), left anterior insula (red), and right anterior insula (orange). Table S1 lists the MNI coordinates of each seed.

#### Seed inter-network connectivity

We quantified the inter-network connectivity strength for a pair of networks. First, the average time course across all seed regions in the first network was correlated with the average time course across all seed regions in the second network. Then, the resulting correlation was Fisher transformed to obtain the inter-network connectivity strength.

#### Head Motion

Since head motion can potentially confound functional connectivity, we included several steps to ensure adequate control of motion confounds. For fetuses, we strictly censored all data for motion and data quality. There were no group differences in motion between the two fetal timepoints. For infants, the mean frame-to-frame displacement was calculated for each run for every individual. Runs with a mean frame-to-frame displacement of more than 0.2 mm were removed from further analysis. Additionally, iterative smoothing and regression of 24 motion parameters (six rigid-body parameters, six temporal derivatives of these parameters, and these 12 parameters squared) were used in the infant data.

### Statistical Analyses

The quantities of interest included within-network connectivity strengths, between-network connectivity strengths, and differences between two such connectivity strengths. To investigate longitudinal changes in such a quantity as a function of PMA, we used a Bayesian spline model^53^ allowing a flexible range of nonlinear behaviors. The mean was modeled using a B-spline basis having a knot at each week of PMA from 30 weeks through 47 weeks. The extent of nonlinearity of the fit was controlled using a hierarchical model incorporating a Gaussian prior distribution on the second-order differences in the spline coefficients and allowing the model to estimate the standard deviation of that Gaussian prior. Random intercepts and slopes were included to account for the repeated measurements of individuals at different gestational ages. The model also allowed the dHCP cohort to have an additive offset and error variance.

Gaussian priors were used for regression coefficients, and half-Cauchy distributions were used for standard deviations, with scales chosen so that the priors were nearly flat for plausible values of the parameters. Posterior probabilities (that is, probabilities conditional on the observed data) were estimated using Markov chain Monte Carlo (MCMC), implemented using JAGS^54^ and R^55^. For each quantity analyzed, the estimated parameters included mean values at the four PMA’s 32, 36, 40, and 44 weeks, changes in the mean from one age to the next ((PMA=36)-(PMA=32), (PMA=40)-(PMA=36), and (PMA=44)-(PMA=40)), as well as the change in mean over the whole range from PMA=32 to PMA=44 weeks. For each such parameter, we report MCMC estimates for the posterior mean (denoted by “m” in the results section), 2.5 and 97.5 percentiles (endpoints of 95% probability credible intervals, denoted by “lower” and “upper”), and the probability of a sign error (denoted by “pr”). The probability of a sign error is the posterior probability that the actual parameter has the opposite sign (negative or positive) from its reported estimated mean^56^. As the probability of a sign error is one-sided, we report statistical “significance” when the probability of a sign error is estimated to be less than 0.05/2=0.025. To investigate the relationship between maternal mental health exposure and functional connectivity, we used the Bayesian spline model modified to include an additional linear effect of the first principal component, PC1, derived from the four maternal prenatal mental health variables.

### Data and code availability

The *Fetal-neonatal Cohort* and *Neonatal Cohort* datasets will be released on https://nda.nih.gov/ following an embargo period. The dHCP data can be accessed at http://www.developingconnectome.org/project/. The image analysis software (BioImage Suite) can be found at https://medicine.yale.edu/bioimaging/suite/ and https://bioimagesuiteweb.github.io/webapp/index.html.

## RESULTS

### Demographic information

Demographics for the subjects are shown in Table 1. Forty-one (49%) were females, the mean birth weight was 3536 (497) g, and the mean PMA at birth was 39.6 (1.3) weeks. There were no significant differences in sex, birth weights, race, ethnicity, or years of maternal education for the Yale cohorts. There were also no differences between the Yale and the dHCP cohorts in sex, birth weight, and PMA. Infants in the Yale longitudinal cohort underwent their first scan (F1) at a mean of 31.2 (0.7) weeks PMA; the second scan (F2) occurred at a mean PMA of 35.3 (0.8) weeks. Although there was no significant difference in PMA at birth for the three cohorts, infants in the dHCP underwent their neonatal scan earlier than those in the Yale cohorts (p<.001).

### Resting-state functional connectivity

#### Intra-network analyses

Longitudinal contrasts for all networks (Tables 2, 3, and 4, Figure 2) revealed significant increases in connectivity strength across the third trimester and perinatal transition (PMA44 – PMA32) as indexed by the mean change in within-network strength between 32 and 44 PMA timepoints: DMN: m=0.199, ECN: m=0.300, and SAL: m=0.289 (pr<0.001 for all three networks). At the network level, all three evidenced significant increases during the prenatal to neonatal transition from 36 to 40 weeks PMA (DMN: 0.073, pr=.002; ECN: 0.161, pr<.001; SAL: 0.161, pr=.012) and continued increases in connectivity during the first postnatal month (40 – 44 weeks; DMN: 0.083, pr=.001; ECN: 0.131, pr=.005; SAL: 0.121, pr=.024).

**Table 2.**
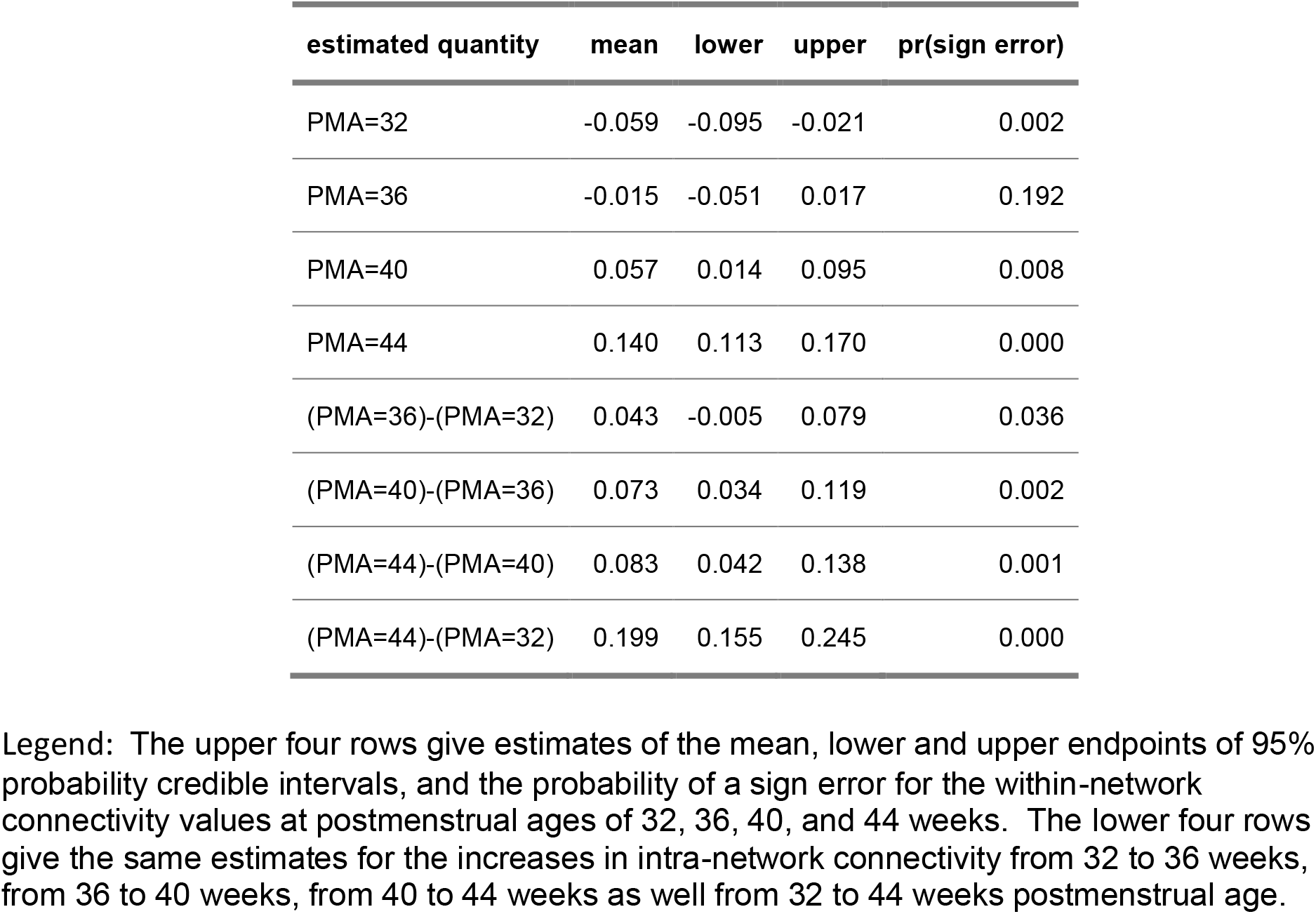
Default Mode Network: Development of intra-network resting functional state connectivity from 32 to 44 postmenstrual weeks

**Table 3.** Executive Control Network: Development of intra-network resting functional state connectivity from 32 to 44 postmenstrual weeks

| **estimated quantity** | **mean** | **lower** | **upper** | **pr(sign error)** |
| --- | --- | --- | --- | --- |
| PMA=32 | -0.037 | -0.091 | 0.014 | 0.077 |
| PMA=36 | -0.028 | -0.074 | 0.017 | 0.110 |
| PMA=40 | 0.133 | 0.062 | 0.210 | 0.001 |
| PMA=44 | 0.264 | 0.225 | 0.303 | 0.000 |
| (PMA=36)-(PMA=32) | 0.009 | -0.053 | 0.071 | 0.391 |
| (PMA=40)-(PMA=36) | 0.161 | 0.082 | 0.252 | 0.000 |
| (PMA=44)-(PMA=40) | 0.131 | 0.039 | 0.217 | 0.005 |
| (PMA=44)-(PMA=32) | 0.300 | 0.240 | 0.365 | 0.000 |

Legend: The upper four rows give estimates of the mean, lower and upper endpoints of 95% probability credible intervals, and the probability of a sign error for the within-network connectivity values at postmenstrual ages of 32, 36, 40, and 44 weeks. The lower four rows give the same estimates for the increases in intra-network connectivity from 32 to 36 weeks, from 36 to 40 weeks, from 40 to 44 weeks as well from 32 to 44 weeks postmenstrual age.

**Table 4.** Salience network: Development of intra-network resting functional state connectivity from 32 to 44 postmenstrual weeks

| **estimated quantity** | **mean** | **lower** | **upper** | **pr(sign error)** |
| --- | --- | --- | --- | --- |
| PMA=32 | -0.006 | -0.072 | 0.070 | 0.418 |
| PMA=36 | 0.001 | -0.060 | 0.069 | 0.495 |
| PMA=40 | 0.163 | 0.059 | 0.264 | 0.002 |
| PMA=44 | 0.283 | 0.233 | 0.331 | 0.000 |
| (PMA=36)-(PMA=32) | 0.007 | -0.081 | 0.094 | 0.430 |
| (PMA=40)-(PMA=36) | 0.161 | 0.027 | 0.284 | 0.012 |
| (PMA=44)-(PMA=40) | 0.121 | 0.001 | 0.236 | 0.024 |
| (PMA=44)-(PMA=32) | 0.289 | 0.196 | 0.370 | 0.000 |

Legend: The upper four rows give estimates of the mean, lower and upper endpoints of 95% probability credible intervals, and the probability of a sign error for the within-network connectivity values at postmenstrual ages of 32, 36, 40, and 44 weeks. The lower four rows give the same estimates for the increases in intra-network connectivity from 32 to 36 weeks, from 36 to 40 weeks, from 40 to 44 weeks as well from 32 to 44 weeks postmenstrual age.

**Figure 2:**
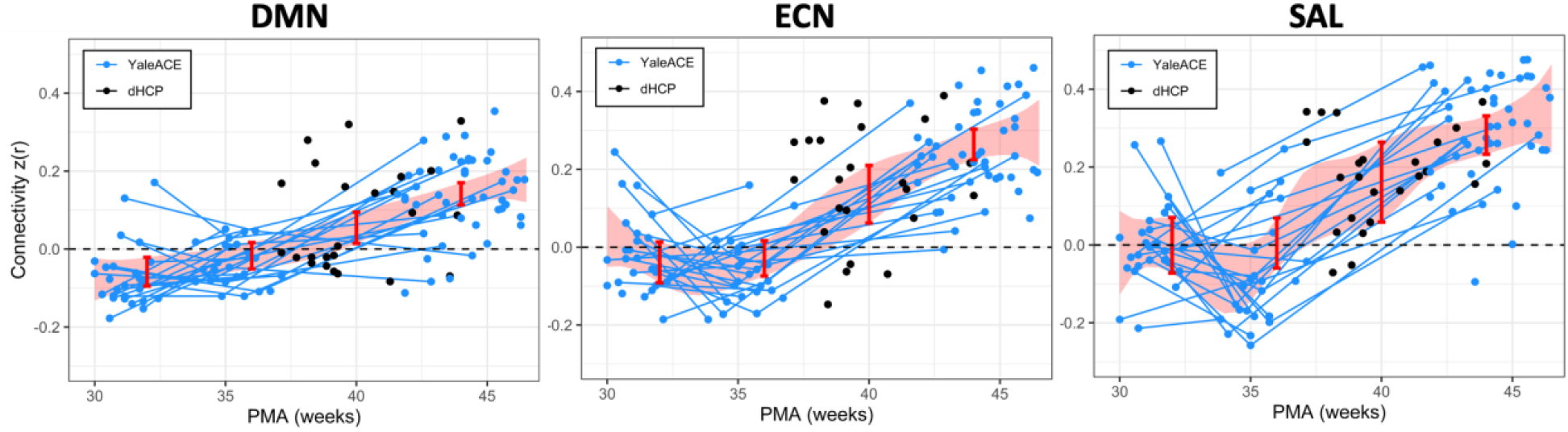
Maturation of intra-network connectivity for the DMN, ECN, and SAL over the third trimester and first postnatal month. Estimates of connectivity strength at the anchors (32, 36, 40, and 44 weeks PMA) for the piecewise linear growth curve are shown as the red bars. The shaded red area represents 95% confidence intervals. Lines indicate longitudinal data from the same participant scanned at multiple time points.

A complementary analysis comparing network strength to zero showed that during the prenatal period, none of the overall network strengths for the DMN, ECN, and Salience networks was significantly greater than 0 at PMA 32 weeks and 36 weeks, while all three networks reached levels significantly greater than 0 levels at 40 weeks PMA (DMN: 0.057, pr=.008; ECN: 0.133, pr=.001; SAL: 0.163, pr=.002; Tables 2, 3 and 4).

In a secondary analysis, we tested the hypothesis that DMN connectivity increased less across the third trimester and perinatal transition than for the ECN or SAL. As shown in Tables S2 – S4, connectivity change across 32 to 44 weeks PMA was significantly less for the DMN compared to SAL (DMN – SAL, PMA44 – PMA32: m=-0.103, pr=0.015). The connectivity change across this time interval was also significantly less for the DMN than ECN (DMN – ECN, PMA44 – PMA32: m=-0.098, pr=.002). In contrast, there was no significant difference in growth curves when the ECN and SAL were compared (SAL – ECN, PMA44 – PMA32: m=0.024, pr=.255).

#### Inter-network analyses

Longitudinal contrasts for inter-network connectivity (Tables 5, 6 and 7; Figure 3) revealed increases in DMN – ECN connectivity strength across the 3^rd^ trimester and perinatal transition (PMA 36 – PMA 32: m=0.029, pr=.037; PMA 40 – PMA 36: 0.042, pr=.027; PMA 44-40 0.066; pr<.001; PMA44 - PMA32: 0.136; pr<.001). In contrast, there were significant decreases in inter-network connectivity for the DMN – SAL and ECN – SAL across this time interval (pr=.004 and pr=.001, respectively).

**Table 5.** Development of inter-network resting state functional connectivity between Default Mode and Executive Control Networks from 32 to 44 postmenstrual weeks.

| **estimated quantity** | **mean** | **lower** | **upper** | **pr(sign error)** |
| --- | --- | --- | --- | --- |
| PMA=32 | 0.024 | -0.001 | 0.050 | 0.031 |
| PMA=36 | 0.053 | 0.029 | 0.076 | 0.000 |
| PMA=40 | 0.095 | 0.053 | 0.124 | 0.000 |
| PMA=44 | 0.161 | 0.141 | 0.183 | 0.000 |
| (PMA=36)-(PMA=32) | 0.029 | -0.003 | 0.056 | 0.037 |
| (PMA=40)-(PMA=36) | 0.042 | -0.001 | 0.073 | 0.027 |
| (PMA=44)-(PMA=40) | 0.066 | 0.034 | 0.119 | 0.000 |
| (PMA=44)-(PMA=32) | 0.136 | 0.106 | 0.169 | 0.000 |

Legend: The upper four rows give estimates of the mean, lower and upper endpoints of 95% probability credible intervals, and the probability of a sign error for the inter-network connectivity values at post menstrual ages of 32, 36, 40, and 44 weeks. The lower four rows give the same estimates for the increases in inter-network connectivity from 32 to 36 weeks, from 36 to 40 weeks, from 40 to 44 weeks as well from 32 to 44 weeks.

**Table 6.**
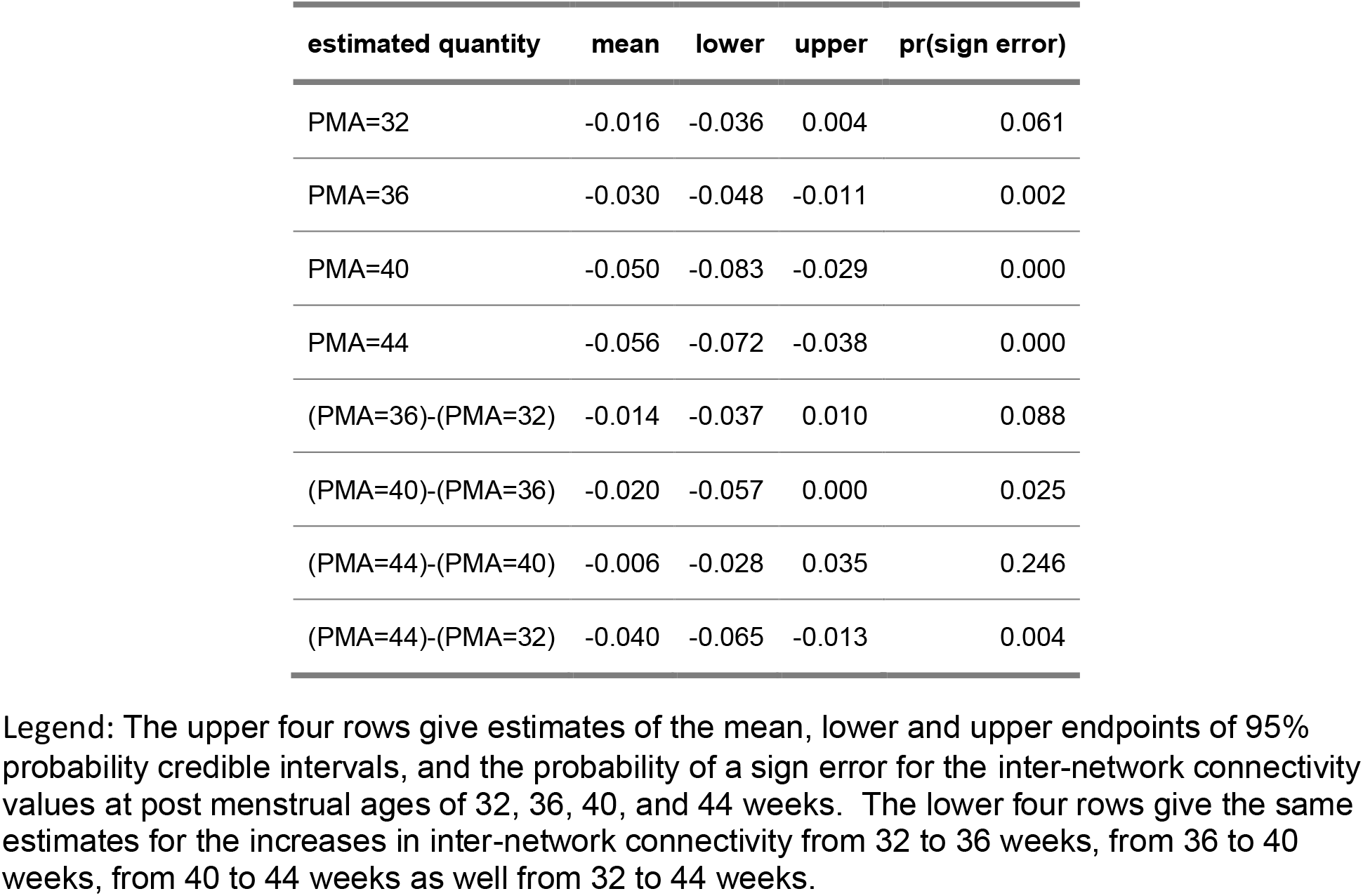
Development of inter-network resting state functional connectivity between Default Mode and Salience Networks from 32 to 44 postmenstrual weeks.

| **estimated quantity** | **mean** | **lower** | **upper** | **pr(sign error)** |
| --- | --- | --- | --- | --- |
| PMA=32 | -0.016 | -0.036 | 0.004 | 0.061 |
| PMA=36 | -0.030 | -0.048 | -0.011 | 0.002 |
| PMA=40 | -0.050 | -0.083 | -0.029 | 0.000 |
| PMA=44 | -0.056 | -0.072 | -0.038 | 0.000 |
| (PMA=36)-(PMA=32) | -0.014 | -0.037 | 0.010 | 0.088 |
| (PMA=40)-(PMA=36) | -0.020 | -0.057 | 0.000 | 0.025 |
| (PMA=44)-(PMA=40) | -0.006 | -0.028 | 0.035 | 0.246 |
| (PMA=44)-(PMA=32) | -0.040 | -0.065 | -0.013 | 0.004 |

Legend: The upper four rows give estimates of the mean, lower and upper endpoints of 95% probability credible intervals, and the probability of a sign error for the inter-network connectivity values at post menstrual ages of 32, 36, 40, and 44 weeks. The lower four rows give the same estimates for the increases in inter-network connectivity from 32 to 36 weeks, from 36 to 40 weeks, from 40 to 44 weeks as well from 32 to 44 weeks.

**Table 7.**
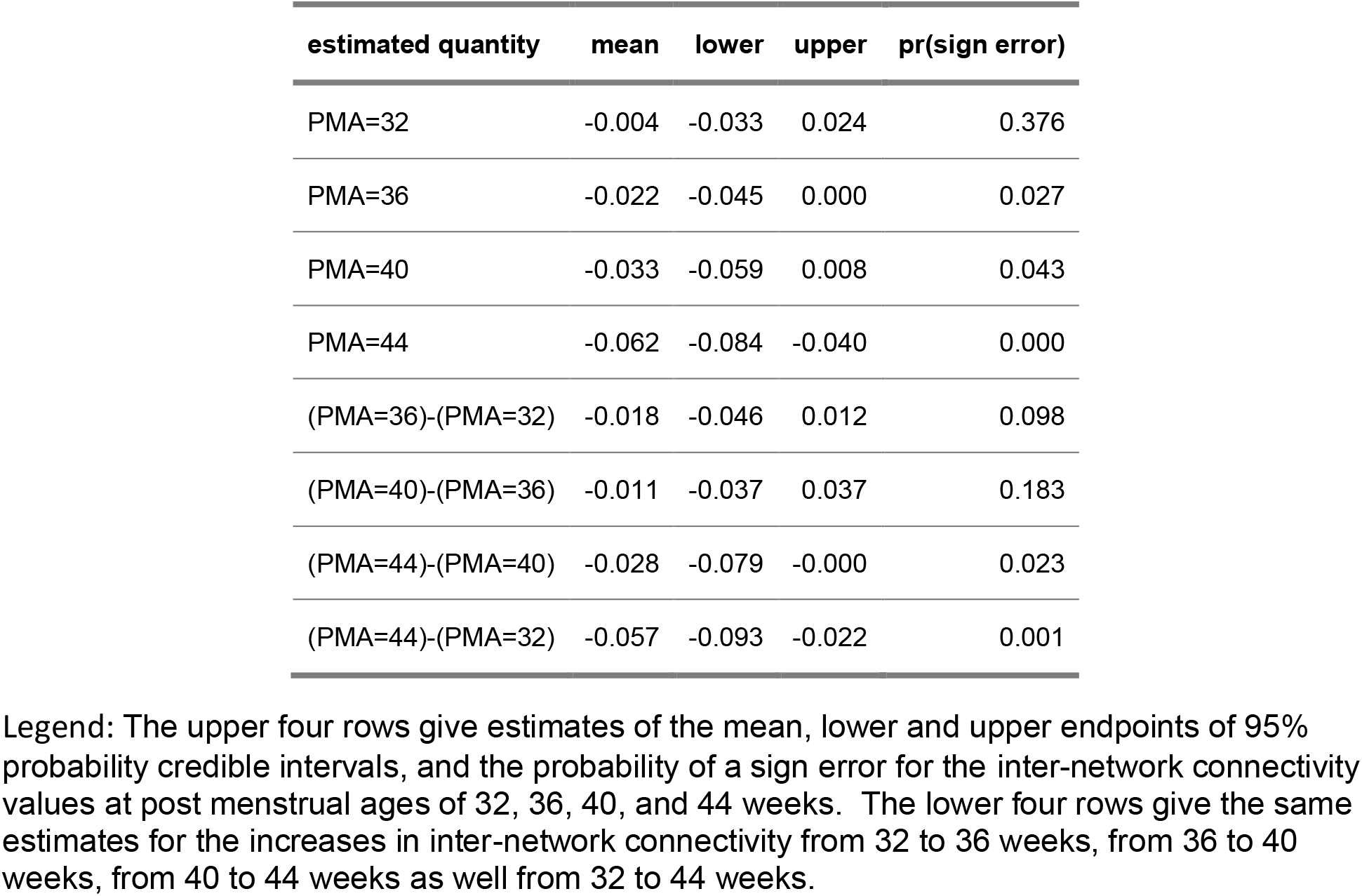
Development of inter-network resting state functional connectivity between Executive Control and Salience Networks from 32 to 44 postmenstrual weeks.

| estimated quantity | mean | lower | upper | pr(sign error) |
| --- | --- | --- | --- | --- |
| PMA=32 | -0.004 | -0.033 | 0.024 | 0.376 |
| PMA=36 | -0.022 | -0.045 | 0.000 | 0.027 |
| PMA=40 | -0.033 | -0.059 | 0.008 | 0.043 |
| PMA=44 | -0.062 | -0.084 | -0.040 | 0.000 |
| (PMA=36)-(PMA=32) | -0.018 | -0.046 | 0.012 | 0.098 |
| (PMA=40)-(PMA=36) | -0.011 | -0.037 | 0.037 | 0.183 |
| (PMA=44)-(PMA=40) | -0.028 | -0.079 | -0.000 | 0.023 |
| (PMA=44)-(PMA=32) | -0.057 | -0.093 | -0.022 | 0.001 |

Legend: The upper four rows give estimates of the mean, lower and upper endpoints of 95% probability credible intervals, and the probability of a sign error for the inter-network connectivity values at post menstrual ages of 32, 36, 40, and 44 weeks. The lower four rows give the same estimates for the increases in inter-network connectivity from 32 to 36 weeks, from 36 to 40 weeks, from 40 to 44 weeks as well from 32 to 44 weeks.

**Figure 3:**
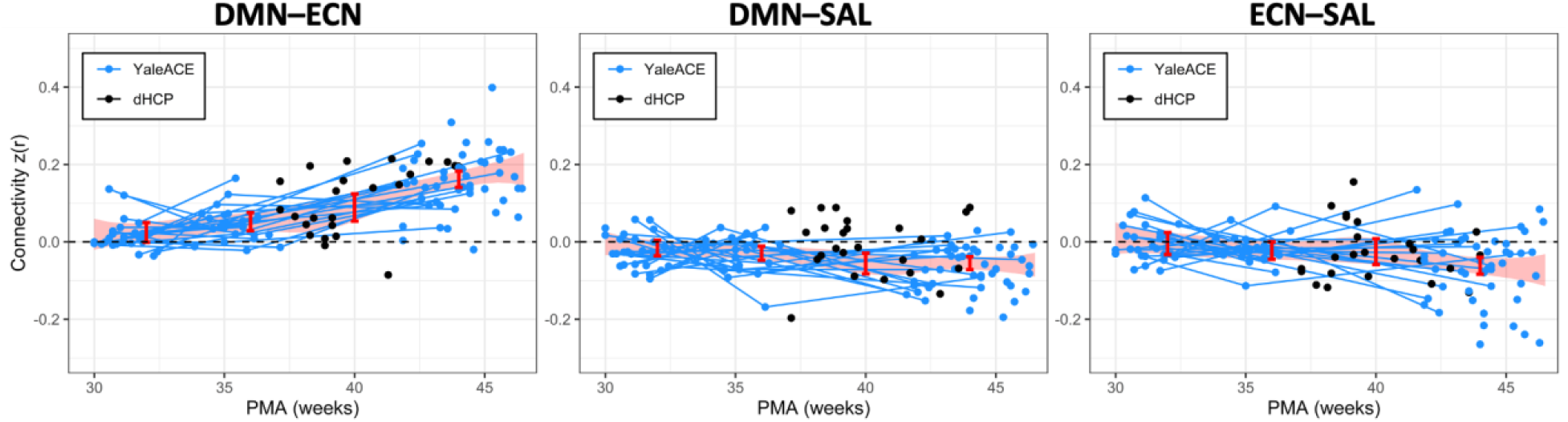
Maturation of inter-network connectivity between DMN–ECN, DMN–SAL, and ECN– SAL over the third trimester and first postnatal month. Estimates of connectivity strength at the anchors (32, 36, 40, and 44 weeks PMA) for the piecewise linear growth curve are shown as the red bars. The shaded red area represents 95% confidence intervals. Lines indicate longitudinal data from the same participant scanned at multiple time points.

As shown in Table 5, the overall inter-network connectivity strength for the DMN - ECN exceeded 0 at 32, 36, 40, and 44 weeks PMA (PMA 32: 0.024, pr=.031; PMA 36: 0.053, pr <.001 PMA 40: 0.095, pr<.001; PMA 44: 0.161, pr<.001). In contrast, the DMN – SAL inter-network analysis showed connectivity significantly < 0 across the 36, 40, and 44 weeks PMA time points (pr≤.002, Table 6). Likewise, for the ECN – SAL inter-network connectivity, the estimate was also significantly < 0 across the 36 – 44 week PMA time interval (p<0.05, Table 7). At 32 weeks, the DMN-SAL and ECN-SAL subtractions were not different from 0 (pr=.061 and pr=.376, respectively).

Longitudinal contrasts for inter-network connectivity (Supplemental Tables 5, 6 and 7) revealed that DMN – ECN connectivity strength increased more than that for DMN – SAL across the 3^rd^ trimester and perinatal transition (PMA 36 – PMA 32: m=0.046, pr=.024; PMA 40 – PMA 36: m=0.064, pr=0.006; PMA 44 – PMA 40: m=0.067, pr=.005; PMA 44 – PMA 32: m=0.177; pr<.001). Likewise, the DMN – ECN connectivity increased more rapidly from PMA 32 weeks to PMA 44 weeks than that for the ECN – SAL inter-network connectivity (PMA 36 – PMA 32: m=0.049, pr=0.033; PMA 40 – PMA 36: 0.050, pr=.076; PMA 44 – PMA 40: 0.0916; pr<.001; PMA 44 – PMA 32: m=0.195; pr<.001). In contrast, there were no significant differences in inter-network connectivity increases between the DMN – SAL and ECN – SAL connections across this time interval.

As shown in Table 5, the difference in the overall inter-network connectivity strength for the DMN - ECN compared to DMN – SAL exceeded 0 across the third trimester and perinatal transition (PMA 32: pr=.012; pr<.001 for all other comparisons). The DMN – ECN inter-network connectivity was also greater than the ECN – SAL connectivity between PMA 36 and PMA 44 (pr≤.002 for all). In contrast, a comparison of connectivity between the ECN – SAL and DMN – SAL showed no values significantly different than zero between PMA 32 and PMA 44 weeks (pr≥.182).

#### Exploratory analysis

*Impact of prenatal mental health on the salience network connectivity* A descriptive summary of the variables that comprise the maternal mental health variable is shown in Table 8. On the STAI screener, 4/29 (13.8%) of mothers exceeded the cutoff of 40 during the prepartum period; 2/29 (6.9%) of the cohort exceeded the cutoff point of 11 on the EPDS, demonstrating that the prevalence of elevated symptoms of depression and anxiety in our sample was consistent with the pre-pandemic estimates observed in the general population.^57-59^

**Table 8.**
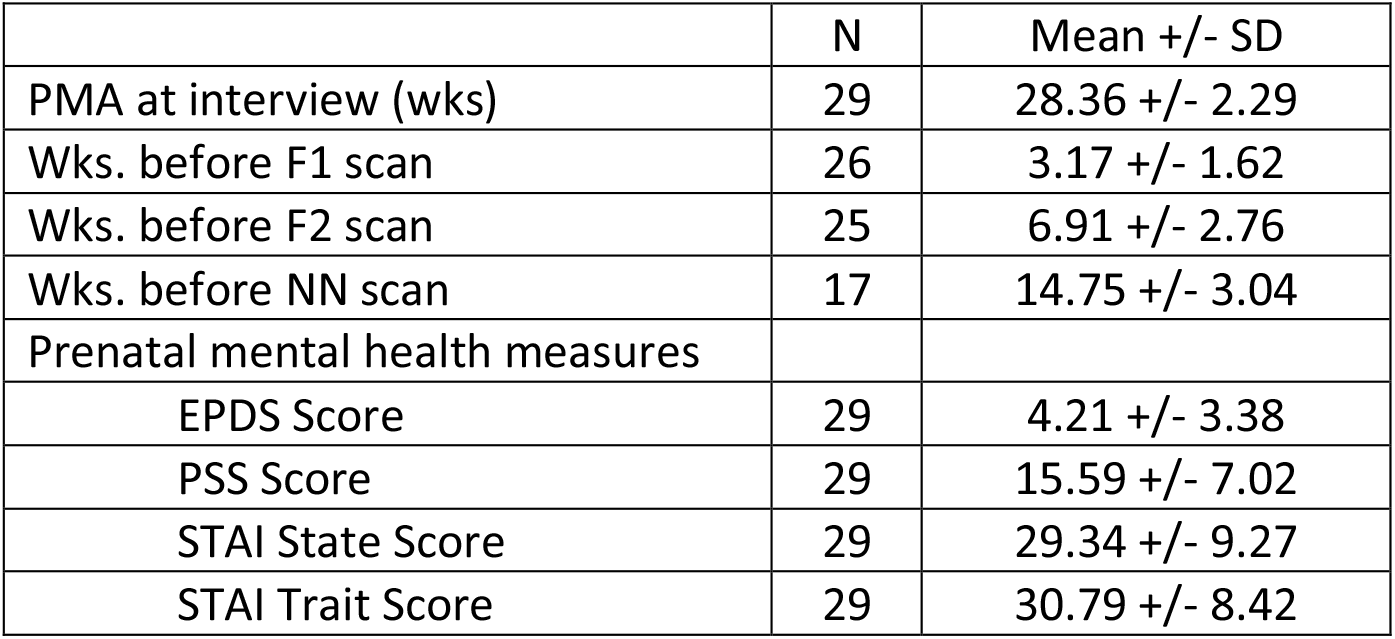
Summary of prenatal mental health measures for the fetal-neonatal cohort

|  | N | Mean +/- SD |
| --- | --- | --- |
| PMA at interview (wks) | 29 | 28.36 +/- 2.29 |
| Wks. before F1 scan | 26 | 3.17 +/- 1.62 |
| Wks. before F2 scan | 25 | 6.91 +/- 2.76 |
| Wks. before NN scan | 17 | 14.75 +/- 3.04 |
| Prenatal mental health measures |  |  |
| EPDS Score | 29 | 4.21 +/- 3.38 |
| PSS Score | 29 | 15.59 +/- 7.02 |
| STAI State Score | 29 | 29.34 +/- 9.27 |
| STAI Trait Score | 29 | 30.79 +/- 8.42 |

Maternal mental health data were collected at a mean PMA of 28.36 (2.29) weeks, or 3.17 (1.62) weeks before the first fetal scan, 6.91 (2.76) weeks prior to the second scan, and 14.75 (3.04) weeks before the neonatal scan. The estimated association between the first principal component PC1 derived from the four maternal prenatal mental health variables and the DM, ECN, and SAL networks was negative for all three networks, with sign error probabilities of 0.079, 0.071, and 0.014, respectively. Thus, while the effects in the ECN and DMN approached significance, only the SAL network had a (one-sided) sign error probability below 0.025. The relationship between PC1 and functional connectivity of the SAL network at the three longitudinal time points is illustrated in Figure 4, which suggests the significant negative impact of maternal mental health across the third trimester and the first postnatal month on the development of the SAL network.

**Figure 4:**
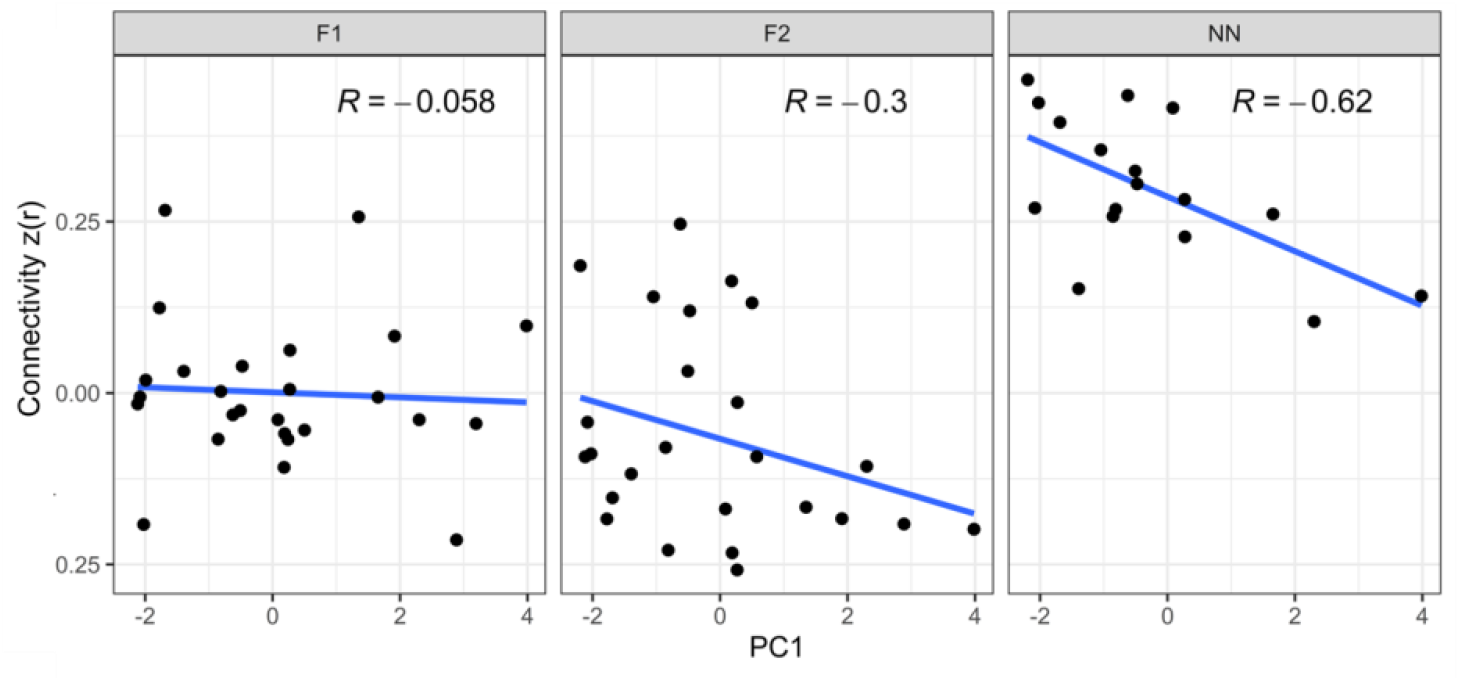
Association between intra-network SAL connectivity at each data collection time point (i.e., F1—early third trimester, F2— mid-third trimester, NN—neonatal) and maternal stress collected in the third trimester. These associations change (possibly strengthening) as intra-network SAL connectivity develops.

## DISCUSSION

The default mode, executive control, and salience networks develop rapidly across the third trimester of gestation and first postnatal month, showing significant changes in inter- and intra-network connectivity across this critical time. At the intra-network level, we report significant increases occurring during the prenatal to neonatal transition for all three networks, with network strength reaching values significantly greater than 0 beginning at 40 weeks PMA for all. These findings suggest that coordination and synchronization between the nodes within the three large cortical networks previously reported in infants^13^ begins before birth, laying foundation for experience-dependent social and cognitive development during postnatal months. Functional connectivity increased less rapidly in the DMN compared to both the ECN and SAL networks, suggesting slower maturation of the network subserving social interactions. Analysis of the connectivity between the three networks revealed both patterns of increasing synchronization and segregation between the networks beginning before birth. Inter-network connectivity greater than 0 for the DMN and ECN was found beginning at 36 weeks and across the first postnatal month, suggesting the emergence of active neural networks in the fetal brain. Finally, in exploratory analyses, we report that higher symptoms of maternal mental health measured at the beginning of the third trimester negatively affected the developmental trajectory of the SAL network across the critical time interval of 30 weeks to 44 weeks PMA.

Inter-network connectivity is critical for engaging the numerous circuits needed for complex behavior.^60^ Neural networks supporting the brain’s ability to process information are present at the beginning of the third trimester,^5^ and intra-network level connectivity has been shown to decrease and increase across the first two years of life in a network specific manner.^13^ Foreshadowing the later data showing increasing integration between the DMN and ECN across the first postnatal year,^13^ our data demonstrate a significant increase in inter-network connectivity strength already during the prenatal period, with a pattern remaining stable from 30 to 44 weeks PMA. This is an intriguing finding,^13^ as later in development these two networks are expected to be anti-correlated at rest.^61 3^ The anticorrelations patterns observed in adults putatively suggest competitive relationship between the networks involved in external attention and internally-focused thought.^11^

Functional connectivity develops from medial to lateral and posterior to anterior.^62,63^ Graph theoretical studies suggest that connectivity between the DMN and ECN during development results from anatomy and their functional development.^60,64,65^ The increasing integration of these from 30 weeks PMA through the first postnatal month may reflect connectivity in the experience-expectant fetal-to-neonatal brain awaiting cognitive demands,^66,67^ focused internal attention,^68^ or a prolonged “transient” local connection.^62,63^

Studies examining the impact of maternal mental health on the development of the DMN, ECN, and SAL have addressed fetuses across the mid-second and third trimesters, preterm neonates, or term infants in the first postnatal month.^27,28,69-75^ A proto-DMN emerges during the third trimester of gestation.^29^ In addition, the anterior insula, the central hub of the SAL, is one of the first cortices to differentiate and develop *in utero*, with insula connectivity reported in the third trimester.^71 76^ Prior fetal studies highlight that maternal distress alters the frontoparietal connectivity of the ECN prior to birth,^29^ and prenatal maternal anxiety influences both ECN and DMN prenatal connections.^26^ Our findings extend the published literature by demonstrating the adverse impact of increasing maternal mental health assessed at the beginning of the third trimester on SAL connectivity across the late third trimester and the first postnatal month.

While exploratory, the associations between prenatal maternal stress and SAL connectivity appear to change and strengthen as the SAL connectivity develops, suggesting that more robust brain-behavioral associations are observable when functional networks are more developed. Alternatively, as maternal stress—and its impact on the developing brain—is likely cumulative, these associations could be dose-dependent. In other words, fetuses experiencing high maternal stress early in the third trimester will likely continue to experience high maternal stress through the neonatal period, leading to more significant differences in brain connectivity. Regardless, these results highlight that the brain correlates of prenatal exposures may change depending on when the imaging and exposure data are collected.^75^

The strengths of this work include the longitudinal/cross-sectional design, robust participant numbers, extensive phenotyping of the fetuses and neonates, collection of multiple standardized maternal health variables early in the third trimester of gestation, and the PMA-appropriate MRI templates. Although follow-up data collection is ongoing, the weaknesses include lacking behavioral data correlating with DMN, ECN, and SAL connectivity. In addition, previously described limitations of fetal functional imaging range from variations in fetal brain orientation, motion, the influence of placental, maternal, and fetal physiological signals, and the small head size to changes in fetal cerebral metabolism and the limited understanding of the physiologic basis of the BOLD fMRI signals in the fetal brain. ^8,47^ These are persisting concerns for MRI studies in neonates and young children, and we have used many of the previously published strategies to address these concerns.^77-79^

We report the emergence of the three networks subserving social cognition in the developing brain across the third trimester and the first postnatal month. The ECN and SAL develop more rapidly than the DMN, but we also show inter-network connectivity greater than 0 between the DMN and ECN beginning at 36 weeks PMA. These fetal networks are plastic and responsive to their environment, and in exploratory analyses, we demonstrate the impact of maternal mental health on fetal network level connectivity. Abnormalities of intra- and inter-network connectivity are becoming well recognized as neuroimaging biomarkers of childhood neurobehavioral disorders.^36^ Future work should target the developmental timing of these abnormalities, identify the genes that support them, and plan early fetal intervention for connectivity disorders of developing brain.^80-82^

## ACKNOWLEDGEMENTS

Research reported in this publication was supported by the National Institute of Mental Health of the National Institutes of Health under Award Number P50MH115716. The content is solely the responsibility of the authors and does not necessarily represent the official views of the National Institutes of Health. The authors thank the families for their participation. We are grateful to Hedy Sarofin, RTR, MR, ARRT, the ACE research fellows Hannah Feiner BS, Rachel Foster BS, Hannah Neiderman BA, and Carolyn Gershman BA for technical assistance, Sahand Negahban, PhD and the ACE Statistics Core for their input on statistical analysis, Kelly Powell PhD, Suzanne Macari PhD, Katherine Kohari MD, Soo Kwon MD, Chelsea Morgan PsyD, James McPartland PhD and Fred Volkmar MD, and Amy Giguere Carney, MSW for their contribution to sample recruitment and characterization.

