## Supplementary material for "Developmental trajectories of the default mode, executive control, and salience networks from the third trimester through the newborn period": SI

### Table S1 Network node coordinates

| Network | ROI | Volume (mm^3) | CenterMassMNIx | CenterMassMNIy | CenterMassMNIz |
| --- | --- | --- | --- | --- | --- |
| ECN | LIPS | 3810 | -40 | -58 | 47 |
|  | RIPS | 3839 | 43 | -58 | 48 |
|  | LDLPFC | 3388 | -32 | 27 | 45 |
|  | RDLPFC | 3085 | 33 | 23 | 45 |
| DMN | RAG | 3800 | 48 | -62 | 23 |
|  | LAG | 3948 | -52 | -67 | 20 |
|  | MPFC | 3555 | 5 | 52 | 9 |
|  | PCC | 2824 | -4 | -40 | 34 |
| SAL | LAINS | 1701 | -41 | 19 | -5 |
|  | RAINS | 1965 | 44 | 17 | -2 |
|  | DACC | 2243 | 1 | 13 | 37 |

Salience Network:

1 : LAINS   (left anterior insula)

2 : RAINS   (right anterior insula)

3 : DACC   (dorsal anterior cingulate)

ECN Network:

1 : LIPS    (left inferior parietal sulcus)

2 : RIPS    (right inferior parietal sulcus)

3 : LDLPFC    (left dorsal lateral pre frontal cortex)

4 : RDLPFC   (right dorsal lateral pre frontal cortex)

Default Mode Network:

1 : RAG    (right angular gyrus)

2 : LAG     (left angular gyrus)

3 : MPFC    (medial prefrontal cortex)

4 : PCC    (posterior cingulate)

### Table S2 Difference between the DMN and SAL trajectories

| **contrasts** | **estimate** | **lower** | **upper** |  |  | **pr(sign error)** |
| --- | --- | --- | --- | --- | --- | --- |
| PMA=32 | -0.041 | -0.111 | 0.026 |  |  | 0.116 |
| PMA=36 | -0.013 | -0.076 | 0.050 |  |  | 0.346 |
| PMA=40 | -0.117 | -0.235 | -0.021 |  |  | 0.011 |
| PMA=44 | -0.144 | -0.196 | -0.086 |  |  | 0.000 |
| (PMA=36)-(PMA=32) | 0.028 | -0.058 | 0.117 |  |  | 0.257 |
| (PMA=40)-(PMA=36) | -0.104 | -0.238 | 0.005 |  |  | 0.029 |
| (PMA=44)-(PMA=40) | -0.027 | -0.139 | 0.120 |  |  | 0.303 |
| (PMA=40)-(PMA=32) | -0.076 | -0.220 | 0.053 |  |  | 0.118 |
| (PMA=44)-(PMA=36) | -0.131 | -0.212 | -0.045 |  |  | 0.002 |
| (PMA=44)-(PMA=32) | -0.103 | -0.186 | -0.012 |  |  | 0.015 |

**Table S3**	Difference between the DMN and ECN trajectories

| **contrasts** | **estimate** | **lower** | **upper** |  |  | **pr(sign error)** |
| --- | --- | --- | --- | --- | --- | --- |
| PMA=32 | -0.018 | -0.073 | 0.043 |  |  | 0.254 |
| PMA=36 | 0.004 | -0.045 | 0.053 |  |  | 0.435 |
| PMA=40 | -0.080 | -0.167 | -0.009 |  |  | 0.014 |
| PMA=44 | -0.116 | -0.158 | -0.073 |  |  | 0.000 |
| (PMA=36)-(PMA=32) | 0.022 | -0.044 | 0.086 |  |  | 0.240 |
| (PMA=40)-(PMA=36) | -0.084 | -0.184 | -0.008 |  |  | 0.016 |
| (PMA=44)-(PMA=40) | -0.036 | -0.121 | 0.069 |  |  | 0.197 |
| (PMA=40)-(PMA=32) | -0.062 | -0.182 | 0.036 |  |  | 0.116 |
| (PMA=44)-(PMA=36) | -0.120 | -0.183 | -0.057 |  |  | 0.000 |
| (PMA=44)-(PMA=32) | -0.098 | -0.166 | -0.035 |  |  | 0.002 |

### Table S4 Difference between the ECN and SAL trajectories

| **contrasts** | **estimate** | **lower** | **upper** |  |  | **pr(sign error)** |
| --- | --- | --- | --- | --- | --- | --- |
| PMA=32 | 0.008 | -0.049 | 0.075 |  |  | 0.400 |
| PMA=36 | 0.011 | -0.041 | 0.061 |  |  | 0.312 |
| PMA=40 | 0.031 | -0.030 | 0.111 |  |  | 0.137 |
| PMA=44 | 0.032 | -0.016 | 0.076 |  |  | 0.084 |
| (PMA=36)-(PMA=32) | 0.003 | -0.073 | 0.064 |  |  | 0.412 |
| (PMA=40)-(PMA=36) | 0.020 | -0.043 | 0.109 |  |  | 0.234 |
| (PMA=44)-(PMA=40) | 0.000 | -0.100 | 0.068 |  |  | 0.413 |
| (PMA=40)-(PMA=32) | 0.023 | -0.070 | 0.119 |  |  | 0.275 |
| (PMA=44)-(PMA=36) | 0.020 | -0.047 | 0.085 |  |  | 0.250 |
| (PMA=44)-(PMA=32) | 0.024 | -0.062 | 0.096 |  |  | 0.255 |

### Table S5 Longitudinal contrasts between DMN – SAL and DMN – ECN inter-network connectivity

| **contrasts** | **estimate** | **lower** | **upper** |  |  | **pr(sign error)** |
| --- | --- | --- | --- | --- | --- | --- |
| PMA=32 | 0.039 | 0.005 | 0.074 |  |  | 0.012 |
| PMA=36 | 0.084 | 0.051 | 0.114 |  |  | 0.000 |
| PMA=40 | 0.149 | 0.107 | 0.186 |  |  | 0.000 |
| PMA=44 | 0.215 | 0.189 | 0.242 |  |  | 0.000 |
| (PMA=36)-(PMA=32) | 0.046 | 0.000 | 0.080 |  |  | 0.024 |
| (PMA=40)-(PMA=36) | 0.064 | 0.025 | 0.109 |  |  | 0.006 |
| (PMA=44)-(PMA=40) | 0.067 | 0.025 | 0.117 |  |  | 0.005 |
| (PMA=40)-(PMA=32) | 0.110 | 0.051 | 0.161 |  |  | 0.002 |
| (PMA=44)-(PMA=36) | 0.131 | 0.094 | 0.174 |  |  | 0.000 |
| (PMA=44)-(PMA=32) | 0.177 | 0.135 | 0.219 |  |  | 0.000 |

### Table S6 Longitudinal contrasts between DMN – ECN and ECN – SAL inter-network connectivity

| **contrasts** | **estimate** | **lower** | **upper** |  |  | **pr(sign error)** |
| --- | --- | --- | --- | --- | --- | --- |
| PMA=32 | 0.028 | -0.018 | 0.074 |  |  | 0.112 |
| PMA=36 | 0.076 | 0.039 | 0.112 |  |  | 0.000 |
| PMA=40 | 0.126 | 0.050 | 0.170 |  |  | 0.002 |
| PMA=44 | 0.222 | 0.189 | 0.259 |  |  | 0.000 |
| (PMA=36)-(PMA=32) | 0.049 | -0.004 | 0.091 |  |  | 0.033 |
| (PMA=40)-(PMA=36) | 0.050 | -0.034 | 0.095 |  |  | 0.076 |
| (PMA=44)-(PMA=40) | 0.096 | 0.046 | 0.193 |  |  | 0.000 |
| (PMA=40)-(PMA=32) | 0.099 | -0.000 | 0.165 |  |  | 0.025 |
| (PMA=44)-(PMA=36) | 0.146 | 0.099 | 0.199 |  |  | 0.000 |
| (PMA=44)-(PMA=32) | 0.195 | 0.139 | 0.254 |  |  | 0.000 |

### Table S7 Longitudinal contrasts between DMN – SAL and ECN – SAL inter-network connectivity

| **contrasts** | **estimate** | **lower** | **upper** |  |  | **pr(sign error)** |
| --- | --- | --- | --- | --- | --- | --- |
| PMA=32 | -0.010 | -0.039 | 0.020 |  |  | 0.228 |
| PMA=36 | -0.008 | -0.032 | 0.019 |  |  | 0.260 |
| PMA=40 | -0.019 | -0.078 | 0.013 |  |  | 0.182 |
| PMA=44 | 0.008 | -0.015 | 0.035 |  |  | 0.275 |
| (PMA=36)-(PMA=32) | 0.003 | -0.031 | 0.037 |  |  | 0.406 |
| (PMA=40)-(PMA=36) | -0.011 | -0.080 | 0.020 |  |  | 0.420 |
| (PMA=44)-(PMA=40) | 0.027 | -0.009 | 0.103 |  |  | 0.108 |
| (PMA=40)-(PMA=32) | -0.009 | -0.081 | 0.035 |  |  | 0.452 |
| (PMA=44)-(PMA=36) | 0.015 | -0.017 | 0.051 |  |  | 0.174 |
| (PMA=44)-(PMA=32) | 0.018 | -0.018 | 0.056 |  |  | 0.161 |
